# The small protein RmpD drives hypermucoviscosity in *Klebsiella pneumoniae*

**DOI:** 10.1101/2020.01.08.899096

**Authors:** Kimberly A. Walker, Logan P. Treat, Virginia L. Miller

**Author notes:** Mailing address: Department of Microbiology and Immunology, University of North Carolina at Chapel Hill, 125 Mason Farm Road, Rm 6101, Campus Box 7290, Chapel Hill, NC 27599-7290. Author Contributions: K.A.W., V.L.M. and L.P.T. designed research; L.P.T. and K.A.W. performed experiments; K.A.W., L.P.T. and V.L.M analyzed data and wrote the paper.

## Abstract

*Klebsiella pneumoniae* has a remarkable ability to cause a wide range of human diseases. It is divided into two broad classes: Classical strains that are a notable problem in healthcare settings due to multidrug resistance, and hypervirulent (*hv*) strains that are drug sensitive, but able to establish disease in immunocompetent hosts. Alarmingly, there has been an increased frequency of clinical isolates that have both drug resistance and *hv*-associated genes. One such gene is *rmpA* that encodes a transcriptional regulator required for maximal capsule (*cps*) gene expression and confers hypermucoviscosity (HMV). This link has resulted in the assumption that HMV is caused by elevated capsule production. However, we recently reported a new *cps* regulator, RmpC, and Δ*rmpC* mutants have reduced *cps* expression but remain HMV, suggesting that capsule and HMV may be separable traits. Here, we report the identification of a small protein, RmpD, that is essential for HMV, but does not impact capsule. RmpD is 58 residues with a putative N-terminal transmembrane domain and highly positively charged C-terminal half, and it is conserved among other *hv K. pneumoniae* strains. Expression of *rmpD in trans* complements both Δ*rmpD* and Δ*rmpA* mutants for HMV, suggesting that RmpD is the key driver of this phenotype. The *rmpD* gene is located between *rmpA* and *rmpC*, within an operon regulated by RmpA. This data, combined with our previous work, suggests a model in which the RmpA-associated phenotypes are largely due to RmpA activating the expression of *rmpD* to produce HMV and *rmpC* to stimulate *cps* expression.

**SIGNIFICANCE:** Capsule is a critical virulence factor in *Klebsiella pneumoniae*, in both antibiotic-resistant classical strains and hypervirulent strains. Hypervirulent strains usually have a hypermucoviscous (HMV) phenotype that contributes to their heightened virulence capacity, but the production of HMV is not understood. The transcriptional regulator RmpA is required for HMV and also activates capsule gene expression, leading to the assumption that HMV is caused by hyperproduction of capsule. We have identified a new gene (*rmpD*) required for HMV but does not contribute to capsule production. This distinction between HMV and capsule production will promote a better understanding of the mechanisms of hypervirulence, which is in great need given the alarming increase in clinical isolates with both drug resistance and hypervirulence traits.

## INTRODUCTION

*Klebsiella pneumoniae* has classically been considered an opportunistic pathogen associated with infection of immunocompromised patients in nosocomial settings (1, 2). Most infections are caused by classical *K. pneumoniae* (cKp) strains and present as pneumonias or urinary tract infections, sometimes leading to bacteremia and septic shock. The widespread occurrence of extended-spectrum β-lactam resistant and carbapenem resistant strains has led both the CDC and WHO to categorize *K. pneumoniae* at the highest level of concern for antibiotic resistant threats (3, 4). In addition, colistin and tigecycline resistant strains of *K. pneumoniae* have been isolated, severely limiting treatment options (5). The deadly case of a pan-resistant cKp strain (resistant to 26 antibiotics) underscores the immense challenge of treating *Klebsiella* infections (6).

In contrast to cKp, the hypervirulent *K. pneumoniae* (hvKp) are community-acquired by immunocompetent individuals (7). HvKp pathology is more severe than that typical of cKp and can include pyogenic liver abscesses, necrotizing fasciitis, meningitis, and endophthalmitis (8, 9). Of particular concern is the emergence of strains with both hypervirulent (*hv*)-associated genes or traits and the multi-drug resistance that is characteristic of cKp (10). Antibiotic resistance genes are often encoded on plasmids (11, 12) and many of the genes corresponding to hypervirulence are carried on large virulence plasmids or mobile genetic elements incorporated on the chromosome (13-15). That these genetic entities can be horizontally transferred suggests there is an increased risk of strains acquiring both hypervirulence and multidrug resistance (16, 17). Alarmingly, there have been recent reports of extensively resistant hypervirulent *K. pneumoniae* (18, 19), and multiple strains where both *hv*-associated genes and antimicrobial resistance genes were present on the same mobile vector have been documented (20-23). These reports of convergence of hypervirulence and antimicrobial resistance in the same strain have heightened the need to better understand how hypervirulence genes interface with a strain’ s genetic background to confer hypervirulent phenotypes. This is particularly important given the extensive genetic diversity of genetic content between *K. pneumoniae* strains.

*Klebsiella* virulence is largely attributable to LPS, pili, a polysaccharide capsule, and siderophores, and these are present in virtually all pathogenic strains (2), heavy metal resistance (24) and hypermucoviscosity (HMV) (2, 25, 26). Capsule is also linked to hypervirulence as the majority of hvKp have type K1 or K2 (27), although *hv*-associated traits have been found in strains with other capsule types (28). Compared to cKp, hvKp produce a thick ‘hypercapsule’ that is thought to contribute to the HMV phenotype.

RmpA is a LuxR-like transcriptional regulator frequently encoded on virulence plasmids or on integrative chromosomal elements (ICE*Kp*) and was initially discovered as a regulator of HMV (14, 25). While the strong correlation between the presence of *rmpA* and hypervirulence has made it a key biomarker for hvKp (27, 29), we still know very little about how *rmpA* contributes to HMV and hypervirulence. Previous studies established that loss of *rmpA* decreases capsule (*cps*) gene expression and reduces HMV in commonly used hvKp strains (30, 31). We recently confirmed these *rmpA*-dependent phenotypes in strain KPPR1S (26). We also described another regulator of capsule gene expression, RmpC, which is encoded downstream of *rmpA*; *rmpA* and *rmpC* are cotranscribed from the same promoter that is positively regulated by RmpA (26). Like *rmpA* mutants, the *rmpC* mutant showed reduced *cps* gene expression but, unlike *rmpA* mutants, retained HMV. We further showed that overexpression of *rmpA* in WT or the Δ*rmpA* and Δ*rmpC* mutants increased HMV. However, overexpression of *rmpA* did not restore *cps* expression in the Δ*rmpC* strain, and overexpression of *rmpC* elevated *cps* expression even in the Δ*rmpA* strain (26). These data suggest that: 1) RmpA is an important determinant for HMV but RmpC is not, 2) reduced *cps* expression in the Δ*rmpA* strain is likely a consequence of reduced *rmpC* expression rather than direct regulation by RmpA, and 3) high levels of *cps* expression are not necessary to confer HMV. This latter conclusion stems from the fact that the Δ*rmpC* mutant has reduced *cps* expression but retains the HMV phenotype, and that exogenous expression of *rmpA* in the Δ*rmpC* strain results in elevated HMV without restoring *cps* expression. Importantly, this was the first clear evidence of a separation between capsule expression and HMV and suggests that HMV is not simply a consequence of elevated capsule production.

Furthering this work, we report here the discovery of a small protein, RmpD, encoded between *rmpA* and *rmpC* that is essential for HMV. The Δ*rmpD* mutant is non-HMV, has no change in *cps* expression, and produces the same amount of uronic acid (capsule) as the wild type parental strain. This provides additional evidence that HMV and capsule result from distinct processes. Expression of *rmpD* is sufficient to confer HMV to a Δ*rmpA* mutant. It is transcribed by the promoter upstream of *rmpA* and therefore is also regulated by RmpA. Thus, it appears that the loss of HMV and *cps* expression observed in *rmpA* mutants is due to reduced transcription of *rmpD* and *rmpC*, respectively, and that the contribution of RmpA to these phenotypes is as an activator of this operon.

## RESULTS

### RmpD is required for hypermucoviscosity

Having made the observation that hypermucoviscosity (HMV) is not necessarily a consequence of elevated capsule expression from examining the individual Δ*rmpA* and Δ*rmpC* strains with complementation plasmids (26), we took one further step by similarly testing a strain that lacked the region encoding both *rmpA* and *rmpC* (strain Δ*rmpA-C*). We predicted that introduction of pRmpA would restore HMV and that pRmpC would restore *manC* expression. HMV was assessed by measuring the OD_600_ of culture supernatants following low-speed centrifugation and expression of *manC* was monitored using a promoter-GFP reporter (*manC* encodes an enzyme that makes one of the K2 sugar precursors and is located in the *cps* locus). While pRmpC resulted in elevated *manC* levels as expected, pRmpA failed to restore HMV (Fig. 1A, B). However, introduction of a plasmid containing the entire region that was deleted resulted in elevated HMV, and in normal levels of *manC* expression (as previously observed), suggesting an element contained within the intergenic space was necessary for HMV. In examining this region, we noted a predicted ORF in the 375 bp region between the *rmpA* and *rmpC* genes (VK055_5098), but this predicted ORF is encoded on the opposite strand (Fig. 1C). The *rmp* locus encoded on the virulence plasmid of NTUH-K2044 also indicates a predicted ORF downstream of *rmpA* (KP1_p021), but this one is encoded on the same strand. The DNA sequence of these loci is very similar between KPPR1S and NTUH-K2044, and the ORF prediction analysis in Geneious R11 identified an ORF in KPPR1S nearly identical to KP1_p021. Thus, we cloned both predicted ORFs from KPPR1S into pMWO-078, transformed them into KPPR1S, Δ*rmpA*, Δ*rmpC* and Δ*rmpAC* strains, and assayed for HMV (Fig. 1D). Introduction of pORF_5098 did not alter HMV, but the KPPR1S homolog of KP1_p021 (pRmpD) resulted in a hyper-HMV phenotype in all strains, including the Δ*rmpAC* strain. Thus, this gene is required for HMV and was named *rmpD*. The region containing *rmpD* is within the *rmp* operon (Fig. S1). Although there is a predicted ORF of 58 amino acids in the DNA sequence cloned in pRmpD, there remained the possibility that this region encoded a regulatory RNA. To distinguish between these possibilities, we generated a plasmid with a *rmpD-2xFLAG* fusion and were able to detect a FLAG-tagged protein of the predicted size (Fig. 1E), indicating that *rmpD* encodes a protein and not a regulatory RNA.

**Figure 1.**
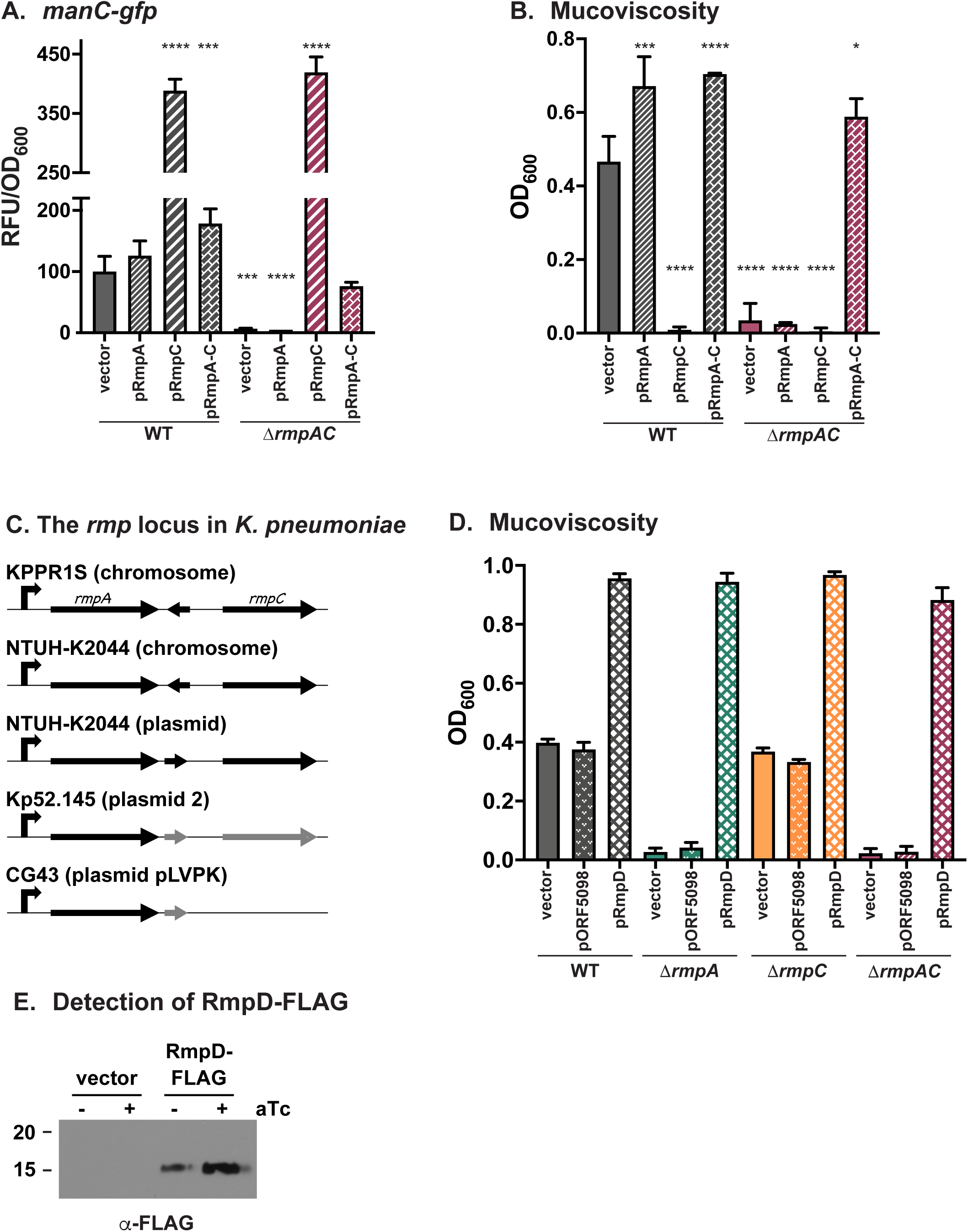
RmpD is required for HMV. Following transformation of the Δ*rmpAC* mutant with pRmpA, pRmpC, or pRmpA-C, *manC* expression (A), and mucosviscosity (B) were assayed as described in Materials and Methods. (C) Schematic of *rmp* loci from several hvKp strains. Black genes, annotated ORFs in NCBI; gray genes, annotated ORFs in Geneious Prime 2019 software. (D) Effect on mucoviscosity of *trans* expression of pORF5098 or pRmpD in WT, Δ*rmpA*, Δ*rmpC* and Δ*rmpA-C* strains. (E) Western blot analysis of whole cell extracts from WT carrying pRmpD-2xFLAG probed with a-FLAG. In (A) and (B), one-way ANOVA with Dunnett’s multiple comparison test was performed using WT with vector as the reference; ****, p ≥ 0.0001; ***, p ≥ 0.001; *, p ≥ 0.05. Data for all but (C) were obtained after 6 h induction of plasmid-borne *rmp* genes.

To further analyze the role of RmpD, we constructed a strain lacking *rmpD* (Δ*rmpD*) and examined *manC* expression and HMV in this mutant. The Δ*rmpD* mutant had wild-type levels of *manC* expression and was non-HMV (Fig. 2). Supporting the notion that it is *rmpD* and not *rmpA* that is necessary for HMV, pRmpA was unable to restore HMV in the Δ*rmpD* mutant. Introduction of pRmpC into the Δ*rmpD* strain resulted in the same high levels of *manC* expression observed in other strains but did not restore HMV. Complementation of Δ*rmpD* with pRmpADC (formerly pRmpA-C) also resulted in an elevated level of HMV. The cultures in which *rmpD* is overexpressed become extremely viscous and have the consistency of a thick syrup (Fig. S2), but have no change in transcription of *manC* (Fig. S3). Collectively, these data suggest that RmpD, rather than RmpA, is necessary for HMV. Given that RmpA regulates the promoter driving expression of *rmpADC* (26), the well-established role of RmpA as a requisite factor for the HMV phenotype is likely due to its function as a transcriptional activator of *rmpD* expression.

**Figure 2.**
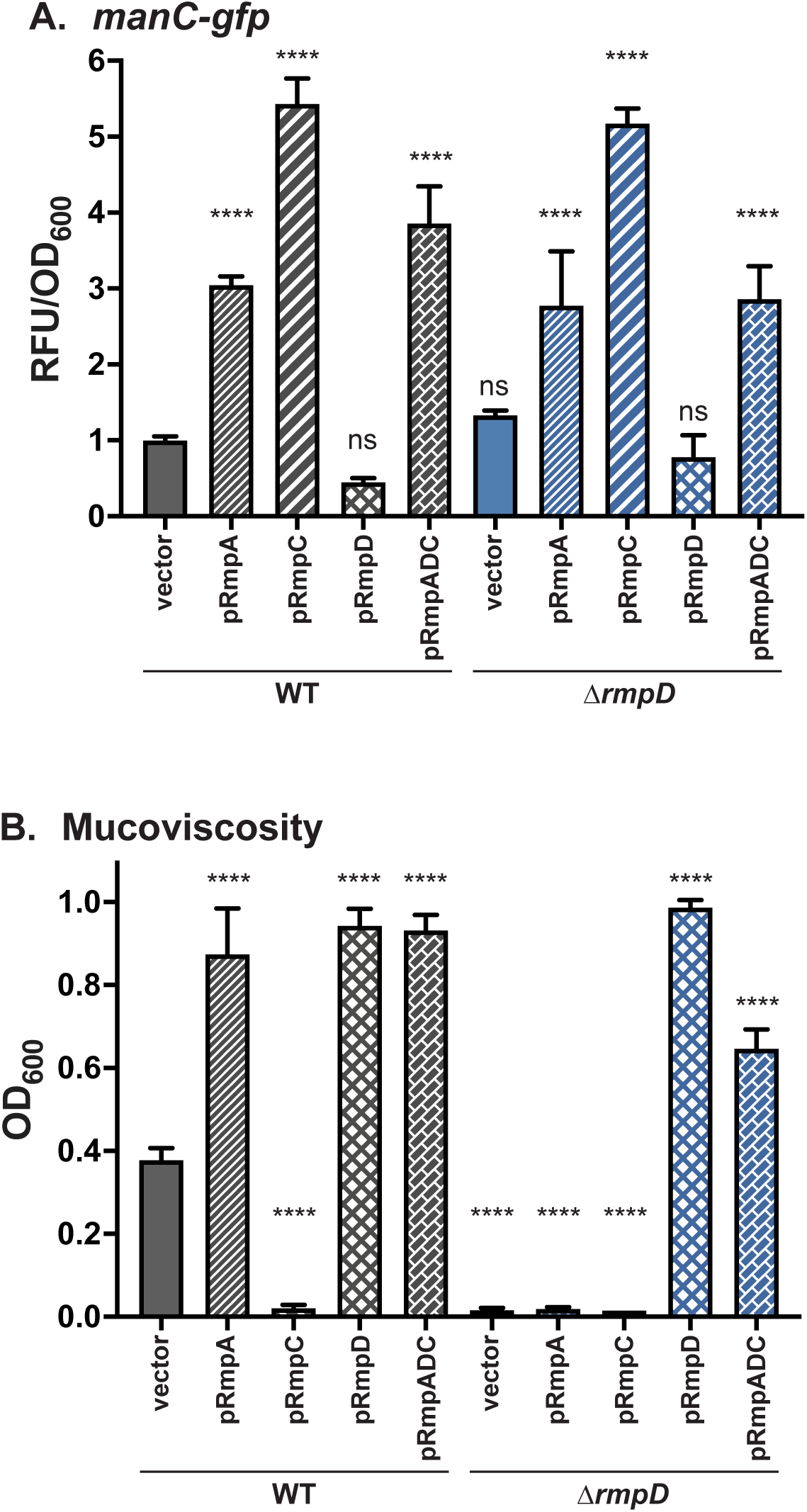
Analysis of Δ*rmpD* strain indicates RmpD but not RmpA is required for HMV. *manC-gfp* expression (A) and mucoviscosity (B) were measured in WT and Δ*rmpD* strains with the indicated plasmids as described in Materials and Methods. Data were obtained after 6 h induction of plasmid-borne *rmp* genes. One-way ANOVA with Dunnett’s post-test was used to determine significance using WT with vector as the reference. ns, not significant; ****, p > 0.0001.

### Impact of *rmpD* in capsule mutants

Contributing to assumptions in the field that HMV is derived from capsule, mutants in hvKp strains that produce no or reduced levels of capsule have also typically been non-HMV (30-33). To further probe the distinction between capsule and HMV, we transformed two capsule mutants (Δ*manC*, Δ*wcaJ*) with pRmpD to determine if these strains could become hyper-HMV. *manC* encodes a GDP-mannose pyrophosphorylase that produces UDP-mannose, one of the sugar precursors of K2 capsule, and *wcaJ* encodes the initiating glycosyltransferase (undecaprenyl phosphotransferase) involved in building the four-sugar K2 subunit. Both capsule mutants, with or without pRmpD, fully sedimented following low speed centrifugation (Fig. 3A), suggesting that the HMV phenotype requires some capsule biosynthetic enzymes and may require capsule production. We therefore examined capsule production in the Δ*rmpD* strain using the uronic acid (UA) assay. Curiously, there was no decrease in UA levels in the Δ*rmpD* strain compared to WT, and addition of pRmpD did not lead to increased UA (Fig. 3B). Collectively, these data imply that production of capsule is not impacted by RmpD, but that at least some components of capsule must be present in order to become HMV. The negative stain, India ink, was used to visualize capsule. WT bacteria show exclusion zones that vary somewhat in size, whereas the *rmpD*-deficient bacteria have thinner, uniform clearing zones (Fig. 4). When *rmpD* is overexpressed in either strain, the bacteria have uniformly large exclusion zones. As predicted, no exclusion zones were observed from staining of Δ*manC* bacteria, although the field has ample bacteria present (Fig. S4). Given that there is no difference in the amount of UA between the WT and Δ*rmpD* strains, these data suggest that the material forming the abundant exclusion zones is different than a typical UA-containing capsule.

**Figure 3.**
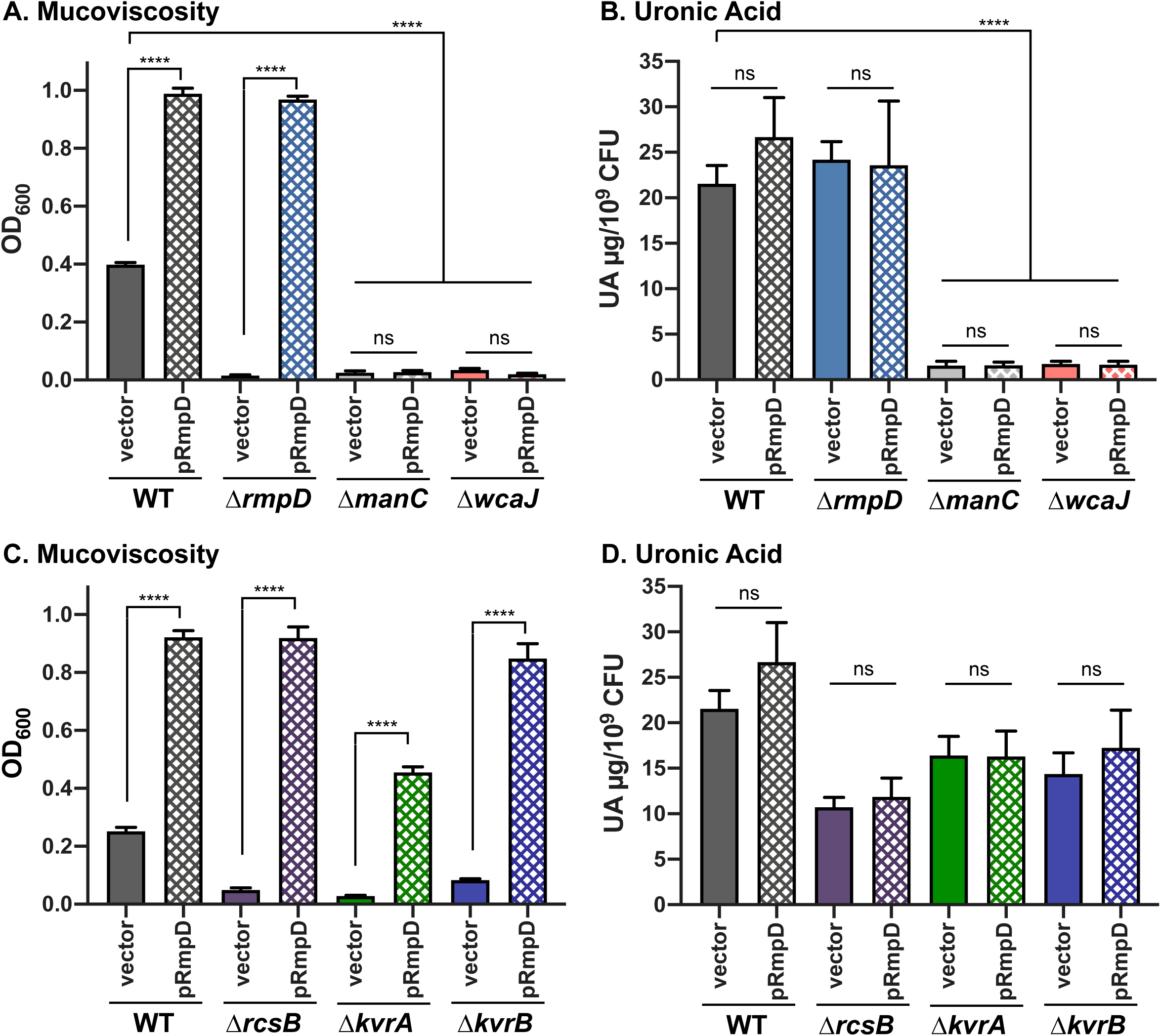
No strong correlation between capsule levels and HMV. Mucoviscosity assay (A) and uronic acid assay (B) of WT, Δ*rmpD*, Δ*manC* and Δ*wcaJ* strains. Mucoviscosity (C) and uronic acid assay (D) assays of WT and regulatory mutants (Δ*rcsB*, Δ*kvrA*, Δ*kvrB*) with vector (pMWO-078) or pRmpD. Data were obtained after 6 h induction of plasmid-borne *rmp* genes as described in Materials and Methods. One-way ANOVA with Tukey’s post-test was used to determine significance to obtain all pairwise comparisons. ns, not significant; ****, p > 0.0001.

**Figure 4.**
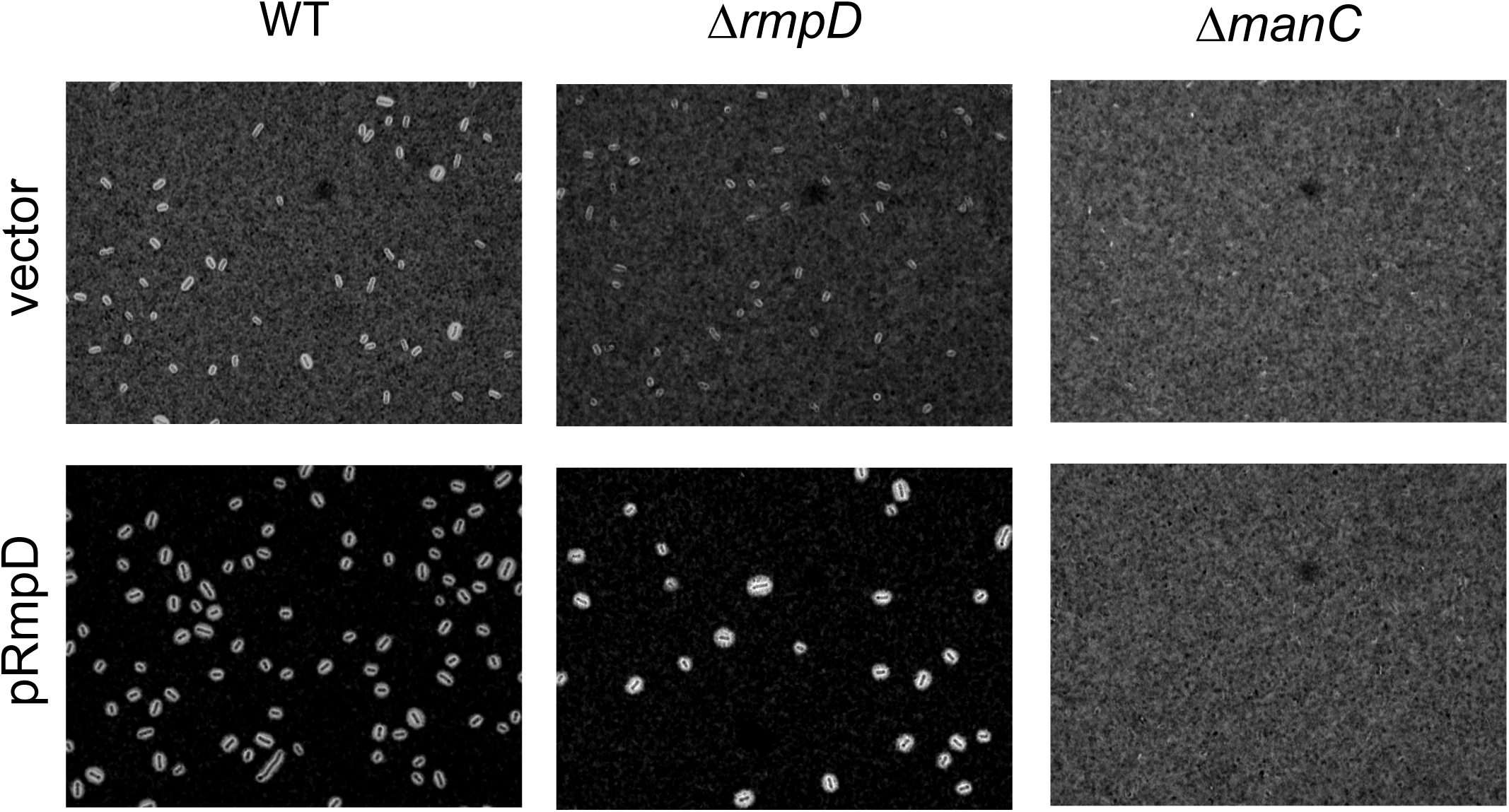
The *rmpD* mutant is encapsulated. Bacteria were stained with India ink and imaged with at 1000x magnification. Capsule is visualized by a clearing zone (ink exclusion) around the bacteria. These strains were expressing gfp; fluorescence images indicating the presence of the bacteria are in Fig. S4. Background shading varies due to uneven liquid distribution under the coverslip.

There are several known regulators of capsule gene expression, of which our lab has identified three and studied five (26, 32). These mutants (Δ*rmpA*, Δ*rmpC*, Δ*kvrA*, Δ*kvrB*, and Δ*rcsB*) all have reduced UA levels and capsule expression, and all but Δ*rmpC* are non-HMV. To further probe the factors necessary for HMV, we transformed the Δ*kvrA*, Δ*kvrB*, and Δ*rcsB* strains with pRmpD and assessed HMV and capsule phenotypes. Each mutant had WT-like HMV, elevated UA levels and elevated *manC* expression with the respective deleted gene complemented in *trans* (Fig. S5). With pRmpD, the Δ*kvrB* and Δ*rcsB* strains became hyper-HMV similarly to the WT strain, and an intermediate level of HMV was observed for the Δ*kvrA* strain (Fig. 3C). Consistent with the results presented above, pRmpD did not restore UA production in these mutants (Fig. 3D), further implying that strains with low capsule expression and UA production are still capable of becoming HMV.

### *rmpD* contributes to immune evasion

One of the virulence phenotypes associated with capsule is the blocking of adherence and phagocytosis (34). To determine if HMV specifically contributed to these processes, we performed adherence assays with the macrophage-like J774A.1 cells. Bacterial strains with pRmpD or vector were grown in the presence of inducer to express *rmpD*, then allowed to interact with J774A.1 cells for 30 minutes. The cells had been pre-treated with cytochalasin D to prevent phagocytosis, allowing measurement of attachment only. After rinsing, the cells were lysed, and each sample was diluted and plated for bacterial enumeration. The WT strain showed about 5% adherence (normalized to inoculum), and the Δ*manC* mutant showed nearly 70% adherence (Fig. 5). The Δ*rmpD* strain behaved like the acapsular *manC* mutant, with ∼85% adherence. The WT or Δ*rmpD* strains with pRmpD were virtually non-adherent, with less than 1% of the bacteria attached. This reduction was not observed in the Δ*manC* mutant with pRmpD, most likely because it remains non-HMV. Because the Δ*rmpD* strain still produces capsule at the level of WT, and the attachment phenotype is the same as a capsule mutant, it appears that the HMV phenotype is the main factor blocking attachment to host cells, and not capsule. Whether capsule has any protective role against adherence, and likely phagocytosis, cannot be fully ascertained from these results, but it is clear that HMV is important as the hyper-HMV strains were almost completely non-adherent.

**Figure 5.**
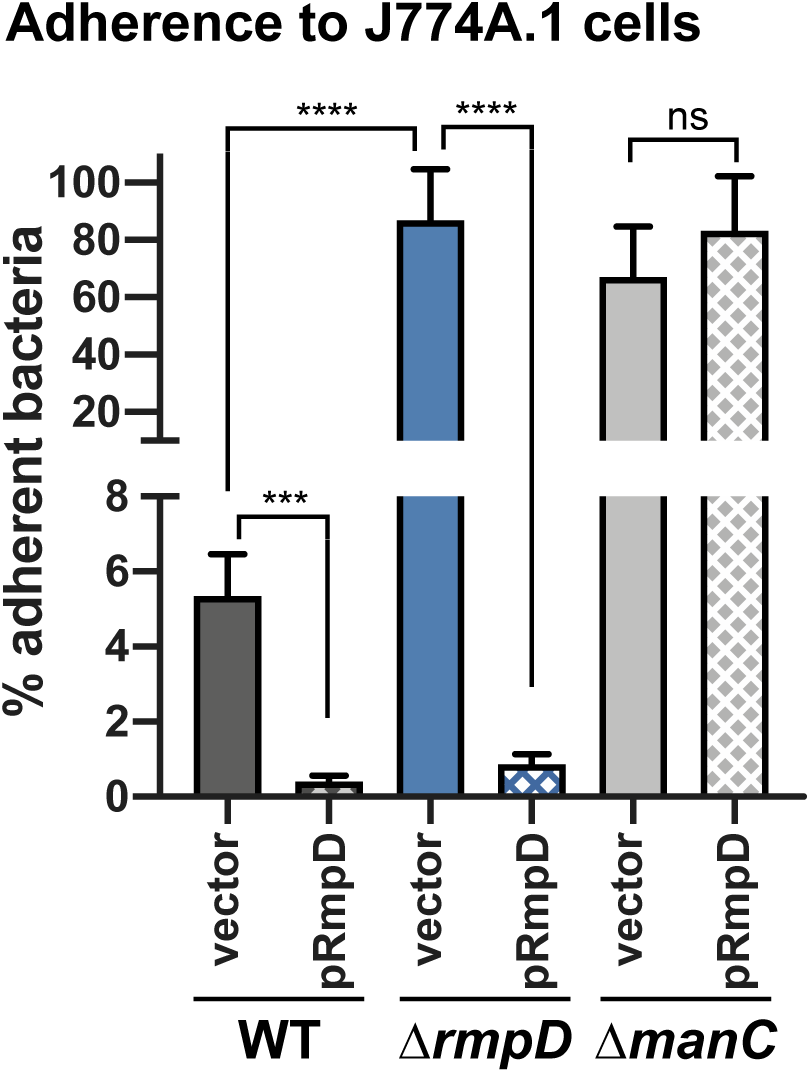
HMV is an anti-adherence factor. J774A.1 cells pretreated with cytochalasin D were inoculated at an MOI of 50, incubated 1 h then rinsed to remove non-adherent bacteria as described in Material and Methods. WT, Δ*rmpD* and Δ*manC* strains carrying either the vector (pMWO-078) or pRmpD were tested. Two-tailed Student’s *t* test was used to determine significance. ns, not significant; ***, p = 0.0001; ****, p > 0.0001.

### Sequence and functional conservation of RmpD homologs

Examination of other hvKp strains reveals that *rmpD* is present when *rmpA* and *rmpC* are present. There are three other hvKp strains that are frequently used for the study of capsule expression and the RmpD proteins in these strains share a high degree of identity (Fig. 6A). To test for functionality, the *rmpD* gene from each of these strains was cloned and expressed in WT and Δ*rmpD* strains of KPPR1S. Not surprisingly, each gene retained the ability to confer hyper-HMV in both WT and Δ*rmpD* strains (Fig. 6B), suggesting that the role of RmpD in HMV is conserved among varying *K. pneumoniae* isolates.

**Figure 6.**
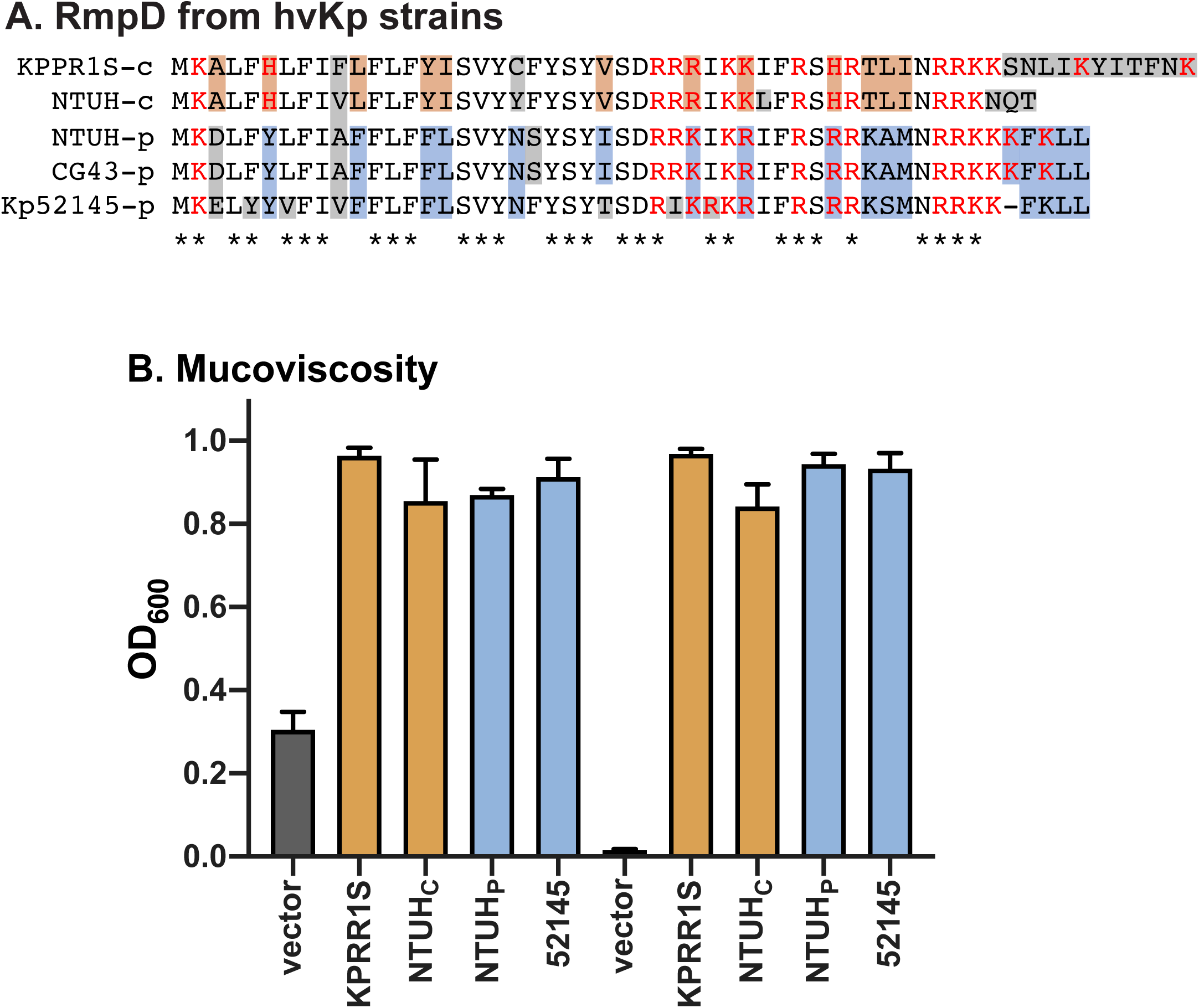
Conservation of *rmpD*. (A) RmpD from known hvKp strains. c, chromosomal copy; p, plasmid copy; orange and blue boxes, residues conserved in chromosomal and plasmid copies, respectively; gray boxes, non-conserved residues; *, fully conserved residues; red residues, positively charged side chains. Accession numbers for these sequences are in the Supplemental Info. (B) Mucoviscosity assay to test if plasmid- and chromosomally-derived *rmpD* genes can complement the Δ*rmpD* mutant for mucoviscosity.

## DISCUSSION

Hypermucoviscosity (HMV) is a phenotype possessed by a subset of *K. pneumoniae* strains and is one of the phenotypes associated with hypervirulent strains (2). RmpA has been established as an essential factor for HMV, and *rmpA* mutants also show reduced capsule gene expression (26, 30, 31). Thus, it has long been assumed that the HMV phenotype was a consequence of abundant capsule production in excess of that observed in classical strains. This arose despite statements in early studies that HMV did not appear to be linked to capsule production (25, 35). However, FITC staining of an *hv K. pneumoniae* strain incubated with K2 antisera suggested the extracapsular substance associated with HMV contained capsular material (36). Although it has been 30 years since the discovery of RmpA, no direct regulation by RmpA of *cps* expression (or other genes) has been demonstrated. In our investigations into the contributions of RmpA to hypervirulence, we confirmed its role in HMV and *cps* expression, but also ascertained that the mechanisms contributing to these phenotypes is much more complex than had been presumed (26). We identified a downstream gene encoding RmpC, a putative transcriptional regulator that modulates *cps* expression, and found that *rmpA* and *rmpC* are in an operon that is autoregulated by RmpA. RmpA and RmpC have overlapping and distinct functions, most notably that the Δ*rmpA* mutant is non-HMV but the Δ*rmpC* mutant retains HMV. Both mutants have a similar reduction in *cps* expression, however, overexpression of *rmpC* complements *cps* expression even in strains lacking *rmpA*. While RmpC has also not been demonstrated to directly regulate *cps* promoters, this data indicated that RmpA was not likely to be a direct regulator of the *cps* genes. We thus concluded that RmpA controlled HMV while RmpC controlled *cps* expression in work that provided the first clear evidence separating the phenotypes of HMV and capsule levels.

In evaluating *cps* expression and HMV in what we thought was a double Δ*rmpA-rmpC* mutant it became clear that the story was not as simple as suggested by the analysis of individual *rmpA* and *rmpC* mutants. Namely, pRmpA did not restore HMV to this Δ*rmpA-C* mutant but a plasmid containing the entire deleted region (pRmpADC) did restore HMV (pRmpC does restore *cps* expression in this mutant). In this current study, we report the initial characterization of RmpD, a small protein encoded in the region between *rmpA* and *rmpC*, also within the *rmp* operon. The data presented here suggest that RmpD is the key factor driving the HMV phenotype. Collectively, our data supports a model in which the role played by RmpA in the HMV and *cps* expression phenotypes is to activate expression of *rmpD* and *rmpC*. This is evidenced by 1) the restoration of HMV in the Δ*rmpA* and Δ*rmpADC* strains with pRmpD, and restoration of *cps* expression in the Δ*rmpA* and Δ*rmpADC* strains with pRmpC, and 2) the inability of pRmpA to restore HMV in the Δ*rmpD* strain or *cps* expression in the Δ*rmpC* strain. Given that several RmpD orthologs were able to complement HMV in the Δ*rmpD* strain, and that *rmpD* is present in strains that also have *rmpA* and *rmpC*, we speculate that RmpD is part of a conserved mechanism conferring HMV to *K. pneumoniae*.

Several lines of evidence further support the notion that production of capsule and HMV are separable. First, deletion of *rmpD* did not alter UA levels, suggesting that production of the capsular material is unaffected by this mutation. Second, strains that are hyper-HMV from overproduction of RmpD did not produce more UA than the WT strain. Third, *trans* expression of *rmpD* in the regulatory mutants (Δ*rmpA*, Δ*kvrA*, Δ*kvrB*, and Δ*rcsB*), that all have reduced *cps* expression and capsule production, were all complemented for HMV. Each of these regulators activate transcription of the *rmpADC* promoter; thus, the loss of HMV in these mutants is most likely due to reduced expression of *rmpD*. Curiously, even though we can detect almost no expression from the *manC* promoter in the Δ*rcsB* strain (26), introduction of pRmpD in the Δ*rcsB* mutant, but not in the Δ*manC* mutant, results in hyper-HMV. Either very low levels of mannose-1-phosphate guanylyltransferase are sufficient for HMV production, or HMV does not actually require this enzyme and the HMV defect in a Δ*manC* strain is an indirect effect of loss of this gene.

In mucosviscosity and adherence assays, the Δ*rmpD* strain behaves nearly identically to the capsule mutant Δ*manC*. Both mutants pellet tightly and are highly adherent to host cells. The hyper-HMV strains (WT and Δ*rmpD* with pRmpD) are essentially non-adherent, but the non-HMV Δ*manC* + pRmpD strain remains highly adherent. This raises the question as to whether or not the anti-adherence property is dependent on capsule or on HMV. Given that the Δ*rmpD* strain is encapsulated, it appears that HMV is a more critical determinant for blocking adherence, and quite likely, in blocking phagocytosis as well. This is consistent with the non-HMV Δ*rmpA* strain having a more severe virulence defect than the HMV-positive Δ*rmpC* strain in the mouse pneumonia model (26). The limited data on the adherence and anti-phagocytic properties of cKp strains makes it difficult to fully extrapolate the significance of capsule in these processes. As HMV has been established as contributing to virulence, *rmpD* mutants are likely to be attenuated in vivo. Support for this comes from re-examination of the virulence defects of the Δ*rmpA* and Δ*rmpC* strains. While it is possible that RmpA regulates additional virulence factors, the loss of *rmpD* expression in the Δ*rmpA* mutant likely contributes to the stronger virulence defect in the Δ*rmpA* mutant than that observed from the Δ*rmpC* mutant. Similarly, analysis of KPPR1 genes essential for infection in a mouse pneumonia model identified mutations in VK055_5096 as deficient for virulence (37). This *orf* is located immediately upstream of *rmpA* (VK055_5097) and the transposon insertion quite likely impaired expression of the *rmp* locus. Furthermore, the virulence plasmid-encoded *rmpD* gene (along with *rmpA* and *rmpC*) was found to be associated with liver abscess formation by NTUH-K2044 (38).

Complicating the notion that HMV is not simply a consequence of overabundant capsule production is that hyper-HMV did not occur in capsule-deficient mutants carrying pRmpD. This suggests that strains can be capsule-positive/HMV-positive or capsule-positive/HMV-negative, but not capsule-negative/HMV-positive. One possible explanation for this is that the HMV material is capsular, but that its export is altered in the presence of RmpD. This situation would mean that even reduced levels of biosynthetic enzymes such as those found in the regulatory mutants are sufficient to yield the extra polysaccharides. A second explanation is that HMV is a polysaccharide distinct from capsule, but that some *cps*-encoded functions are required to produce this material. A third possibility is that the HMV material is a modified form of capsule. The presence of RmpD could influence synthesis or export of the altered polysaccharide. That capsule-like material is part of HMV material is supported by the K2-positive staining of the HMV substance from a WT strain but not from non-HMV mutants (36).

To date, HMV has primarily been associated with *hv* K1 and K2 strains, but more than 130 capsule types of *K. pneumoniae* have been identified (39). Of significant concern is the number of recent reports of strains with both carbapenem resistance and *hv*-associated genes, including *rmpADC*. These strains are genetically quite distinct (including capsule type) from the hvKp that have been circulating, and it is not known to what degree acquisition of the *rmpADC* locus will impact HMV and virulence of these strains. While we have shown that RmpD from either a K2 or K1 strain can confer HMV in a K2 strain, it is not clear if there is capsule type specificity for this RmpD function. We also do not know, beyond a few *cps* genes, what, if any, other genes are necessary to confer HMV or if these genes are conserved in all *K. pneumoniae* strains. A better understanding of what is required for HMV and how genetic background influences the HMV associated hypervirulent phenotypes will be important for determining the risks associated with CR-cKp strains that acquire *rmpADC*.

## MATERIALS and METHODS

More detail can be found in the Supplementary Information

### Bacterial strains, plasmids and growth conditions

The strains and plasmids used in this work are listed in Table S1. *E. coli* strains were grown in LB medium at 37°C. *K. pneumoniae* were grown at 37°C in M9 medium supplemented with 0.4% glucose and 0.2% casamino acids (M9-CAA). Unless otherwise noted, saturated overnight cultures were diluted to OD_600_ = 0.2 and grown for 6 h. Antibiotics were used where appropriate: kanamycin (Kan), 50 μg/ml; rifampicin (Rif), 30 μg/ml, spectinomycin (Sp), 50 μg/ml. For expression of genes cloned into pMWO-078, 100 ng/ml anhydrous tetracycline (aTc) was added to the media at the time of subculture. The primers used for cloning are listed in Table S2. In-frame gene deletions in *K. pneumoniae* were constructed by allelic exchange using pKAS46-based plasmids as described (26).

Complementation plasmids were constructed using pMWO-078 (40). Plasmids containing promoter-*gfp* fusions were cloned in pPROBE-tagless (41). The *gfp* reporter and complementation plasmids were introduced into *K. pneumoniae* by electroporation as described (26).

### Transcriptional *gfp* reporter assays

Relative fluorescent units (RFU) and OD_600_ were measured from bacterial cultures diluted 1:10 using a Synergy H1 plate reader (Bio-Tek, Winooski, WI) and a Bio-Rad spectrophotometer (Bio-Rad, Hercules, CA), respectively. Data are presented as RFU/OD_600_, normalized to the activity from the wild type strain in each assay.

### Assessment of capsule production and HMV

Uronic acid was measured essentially as described (42). Mucoviscosity of liquid cultures was determined by measuring the OD_600_ of the culture supernatant following low-speed centrifugation as described (26).

### Immunoblotting

Whole cell lysates from cultures grown in M9-CAA with aTc for 6 h were separated on 15% SDS-PAGE gels, transferred to PVDF membranes and probed with α-FLAG antibody (Sigma, M2 monoclonal antibody) and detected with chemiluminescence.

### Adherence assays

Adherence assays were performed essentially as described (32) using J774A.1 cells. The cells were pretreated with cytochalasin D 1 h prior to inoculation to prevent internalization of the bacteria. The adherent bacteria (recovered CFU) are reported as a percent of the inoculum CFU.

### India ink staining

Bacterial cultures carrying a constitutively expressing *gfp* reporter (pJH026) were grown as for all other assays. Equal volumes of culture and India ink were mixed on a glass slide and overlaid with a coverslip. Microscopy was performed using a Keyence BZ-X810 microscope at 1000x magnification. Images vary some due to the irregular spreading of the liquid on the slide.

### Statistics and Replicates

Statistical tests for each experiment are given in the figure legends and were performed using GraphPad Prism 8.2. In every assay, a minimum of three assays were performed, each with biological replicates. Typically, a representative experiment is presented.

## ACKNOWLEDGEMENTS

We thank Rita Tamayo for thoughtful discussions on this manuscript and for use of the Keyence BZ-X810 microscope. This work was supported by R21AI132925 to V.L.M. from NIAID.

